# Accurate and Versatile 3D Segmentation of Plant Tissues at Cellular Resolution

**DOI:** 10.1101/2020.01.17.910562

**Authors:** Adrian Wolny, Lorenzo Cerrone, Athul Vijayan, Rachele Tofanelli, Amaya Vilches Barro, Marion Louveaux, Christian Wenzl, Susanne Steigleder, Constantin Pape, Alberto Bailoni, Salva Duran-Nebreda, George Bassel, Jan U. Lohmann, Fred A. Hamprecht, Kay Schneitz, Alexis Maizel, Anna Kreshuk

**Affiliations:** HCI-IWR, Heidelberg University, Germany; EMBL Heidelberg, Germany; School of Life Sciences, Technical University of Munich, Germany; COS, Heidelberg University, Germany; School of Life Sciences, The University of Warwick, UK

**Author notes:** These authors contributed equally to this work (Adrian Wolny, Lorenzo Cerrone).

## Abstract

Quantitative analysis of plant and animal morphogenesis requires accurate segmentation of individual cells in volumetric images of growing organs. In the last years, deep learning has provided robust automated algorithms that approach human performance, with applications to bio-image analysis now starting to emerge. Here, we present PlantSeg, a pipeline for volumetric segmentation of plant tissues into cells. PlantSeg employs a convolutional neural network to predict cell boundaries and graph partitioning to segment cells based on the neural network predictions. PlantSeg was trained on fixed and live plant organs imaged with confocal and light sheet microscopes. PlantSeg delivers accurate results and generalizes well across different tissues, scales, and acquisition settings. We present results of PlantSeg applications in diverse developmental contexts. PlantSeg is free and open-source, with both a command line and a user-friendly graphical interface.

## 1 Introduction

Large-scale quantitative study of morphogenesis in a multicellular organism entails an accurate estimation of the shape of all cells across multiple specimen. State-of-the-art light microscopes allow for such analysis by capturing the anatomy and development of plants and animals in terabytes of high-resolution volumetric images. With such microscopes now in routine use, segmentation of the resulting images has become a major bottleneck in the downstream analysis of large-scale imaging experiments. A few segmentation pipelines have been proposed [1, 2], but these either do not leverage recent developments in the field of computer vision or are difficult to use for non-experts.

With a few notable exceptions, such as the Brainbow experiments [3], imaging cell shape during morphogenesis relies on staining of the plasma membrane with a fluorescent marker. Segmentation of cells is then performed based on their boundary prediction. In the early days of computer vision, boundaries were usually found by edge detection algorithms [4]. More recently, a combination of edge detectors and other image filters was commonly used as input for a machine learning algorithm, trained to detect boundaries [5]. Currently, the most powerful boundary detectors are based on Convolutional Neural Networks (CNNs) [6, 7, 8]. In particular, the U-Net architecture [9] has demonstrated excellent performance on 2D biomedical images and has later been further extended to process volumetric data [10].

Once the boundaries are found, other pixels need to be grouped into objects delineated by the detected boundaries. For noisy, real-world microscopy data, this post-processing step still represents a challenge and has attracted a fair amount of attention from the computer vision community [11, 12, 13, 14, 15]. If centroids (“seeds”) of the objects are known or can be learned, the problem can be solved by the watershed algorithm [16, 17]. For example, in [18] a 3D U-Net was trained to predict cell contours together with cell centroids as seeds for watershed in 3D confocal microscopy images. This method, however, suffers from the usual drawback of the watershed algorithm: misclassification of a single cell centroid results in sub-optimal seeding and leads to segmentation errors.

Recently an approach combining the output of two neural networks and watershed to detect individual cells showed promising results on segmentation of cells in 2D [19]. Although this method can in principle be generalized to 3D images, the necessity to train two separate networks poses additional difficulty for non-experts.

While deep learning-based methods define the state-of-the-art for all image segmentation problems, only a handful of software packages strives to make them accessible to non-expert users in biology (reviewed in [20]). Notably, the U-Net segmentation plugin for ImageJ [21] conveniently exposes U-Net predictions and computes the final segmentation from simple thresholding of the probability maps. CDeep3M [22] and DeepCell [23] enable, via the command-line, the thresholding of the probability maps given by the network, and DeepCell allows instance segmentation as described in [19]. More advanced post-processing methods are provided by the ilastik Multicut workflow [24], however, these are not integrated with CNN-based prediction.

Here, we present PlantSeg, a deep learning-based pipeline for volumetric instance segmentation of dense plant tissues at single-cell resolution. PlantSeg processes the output from the microscope with a CNN to produce an accurate prediction of cell boundaries. Building on the insights from previous work on cell segmentation in electron microscopy volumes of neural tissue [13, 15], the second step of the pipeline delivers an accurate segmentation by solving a graph partitioning problem. We trained PlantSeg on 3D confocal images of fixed *Arabidopsis thaliana* ovules and 3D+t light sheet microscope images of developing lateral roots, two standard imaging modalities in the studies of plant morphogenesis. We investigated a range of network architectures and graph partitioning algorithms and selected the ones which performed best with regard to extensive manually annotated groundtruth. We benchmarked PlantSeg on a variety of datasets covering a range of plant organs and image resolutions. Overall, PlantSeg delivers excellent results on unseen data, and, as we show through quantitative and qualitative evaluation, new datasets do not necessarily require network retraining. PlantSeg is a free and open source tool which contains the complete pipeline for segmenting large volumes. Each step of the pipeline can optionally be adjusted via a convenient graphical user interface while expert users can modify the Python code or provide their own pre-trained networks for the first step of the pipeline. Besides the tool itself, we provide all the networks we trained for the confocal and light sheet modalities at different resolution levels and make all our training and validation data publicly available.

## 2 Results

### 2.1 A pipeline for segmentation of plant tissues into cells

The segmentation algorithm we propose contains two major steps. In the first step, a fully convolutional neural network (a 3D U-Net) is trained to predict cell boundaries. Afterwards, a region adjacency graph is constructed from the pixels with edge weights computed from the boundary predictions. In the second step, the final segmentation is computed as a partitioning of this graph into an unknown number of objects (see Figure 1). Our choice of graph partitioning as the second step is inspired by a body of work on segmentation for nanoscale connectomics (segmentation of cells in electron microscopy images of neural tissue), where such methods have been shown to outperform more simple post-processing of the boundary maps [13, 15, 25].

**Figure 1:**
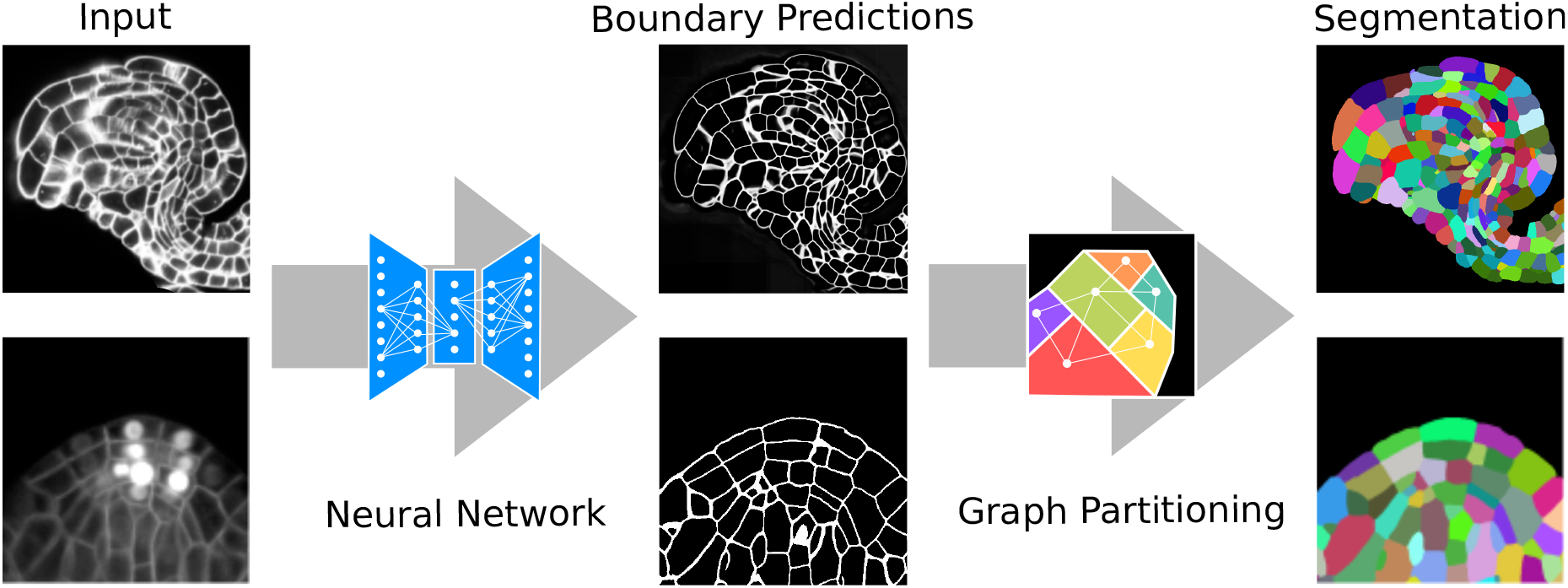
Segmentation of plant tissues into cells using PlantSeg. First, PlantSeg uses a 3D UNet neural network to predict the boundaries between cells. Second, a volume partitioning algorithm is applied to segment each cell based on the predicted boundaries. The neural networks were trained on ovules (top, confocal laser scanning microscopy) and lateral root primordia (bottom, light sheet microscopy) of *Arabidopsis thaliana*.

#### 2.1.1 Datasets

To make our tool as generic as possible, we used both fixed and live samples as core datasets for design, validation and testing. Two microscope modalities common in studies of morphogenesis were employed, followed by manual and semi-automated annotation of groundtruth segmentations.

The first dataset consists of fixed *Arabidopsis thaliana* ovules at all developmental stages acquired by confocal laser scanning microscopy with a voxel size of 0.075×0.075×0.235 *μ*m. 48 volumetric stacks with hand-curated groundtruth segmentation were used. A complete description of the image acquisition settings and the groundtruth creation protocol is reported in [26].

The second dataset consists of three time-lapse videos showing the development of *Arabidopsis thaliana* lateral root primordia (LRP). Each recording was obtained by imaging wild-type Arabidopsis plants expressing markers for the plasma membrane and the nuclei [27] using a light sheet fluorescence microscope (LSFM). Stacks of images were acquired every 30 minutes with constant settings across movies and time points, with a voxel size of 0.1625 × 0.1625 × 0.250 *μ*m. The first movie consists of 52 time points of size 2048 × 1050 × 486 voxels. The second movie consists of 90 time points of size 1940 × 1396 × 403 voxels and the third one of 92 time points of size 2048 × 1195 × 566 voxels. The groundtruth was generated for 27 images depicting different developmental stages of LRP coming from the three movies (see Supp. Appendix A).

The two datasets were acquired on different types of microscopes and differ in image quality. To quantify the differences we used the peak signal-to-noise (PSNR) and the structural similarity index measure (SSIM) [28]. We computed both metrics using the input images and their groundtruth boundary masks; higher values show better quality. The average PSNR measured on the light sheet dataset was 22.5 ± 6.5 dB (average ± SD, *n* = 4), 3.4 dB lower than the average PSNR computed on the confocal dataset (25.9 ± 5.7, *n* = 7). Similarly, the average SSIM is 0.53 ± 0.12 for the light sheet, 0.1 lower than 0.63 ± 0.13 value measured on the confocal images. Both datasets thus contain a significant amount of noise. LSFM images are noisier and more difficult to segment, not only because of the noise, but also due to part of nuclear labels being in the same channel as membrane staining.

In the following we describe in detail the design and performance of each of the two steps of the pipeline.

#### 2.1.2 Step 1: cell boundary detection

We performed quantitative comparison of several design choices in the network architecture, loss function and training protocol. We trained one set of CNNs for each dataset as the ovule and lateral root datasets are substantially different.

In more detail, we compared the regular 3D U-Net as described in [10] with a Residual U-Net from [29]. We tested two loss functions commonly used for the semantic segmentation task: binary cross-entropy 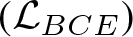 [9] and Dice loss 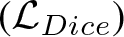 We additionally investigated whether the size of patches used for training influenced the performance. For this, we compared training on a single large patch, versus training on multiple smaller patches per network iteration. For single patch we used group normalization [30] whereas standard batch normalization [31] was used for the multiple patches. Finally, besides the regular data augmentation techniques, we add more training data by corrupting the training volumes by additive Gaussian and Poisson noise with random settings.

In the ovule dataset, 39 stacks were randomly selected for training, two for validation and seven for testing. In the LRP dataset, 27 time points were randomly selected from the three videos for training, two time points were used for validation and four for testing.

The performance of different CNNs is illustrated by the precision/recall curves evaluated at different threshold levels of the predicted boundary probability maps (Figure 2). All models performed well on both the ovule and the LRP datasets (see also Supp. Figure 3 and Supp. Table 2). On the ovule dataset training with several smaller patches and batch normalisation performed better. Noise augmentation was beneficial for both modalities.

**Figure 2:**
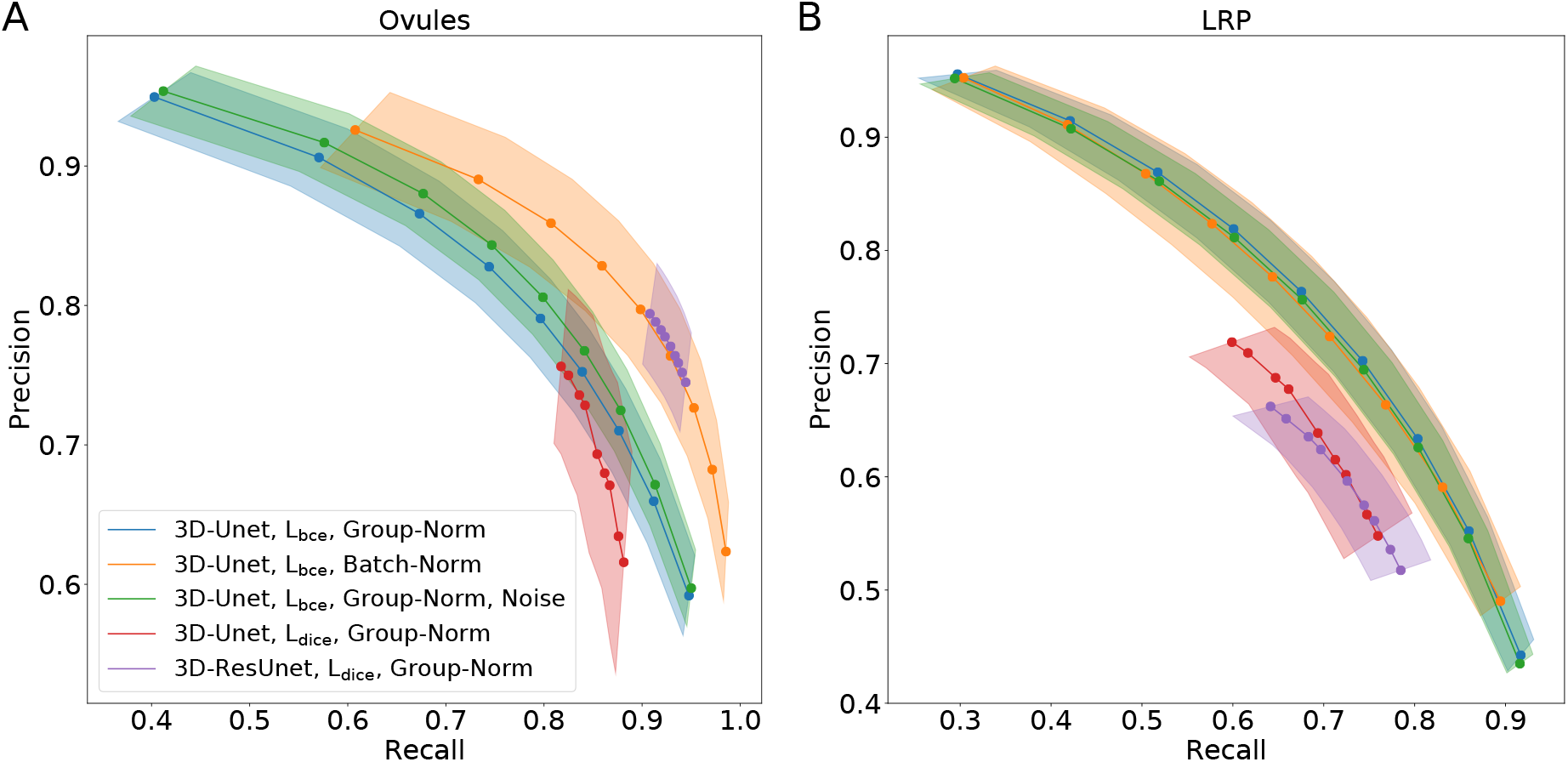
Precision-recall curves for different CNN variants on the ovule (A) and lateral root primordia (B) datasets. Accuracy of boundary prediction was assessed for five training procedures that sample different type of architecture (3D U-Net *vs.* 3D Residual U-Net), loss function (binary cross-entropy *vs.* Dice) and normalization (Group-Norm *vs.* Batch-Norm). Points are average of seven (ovules) and four (LRP) values and the shaded area represent the standard error.

**Figure 3:**
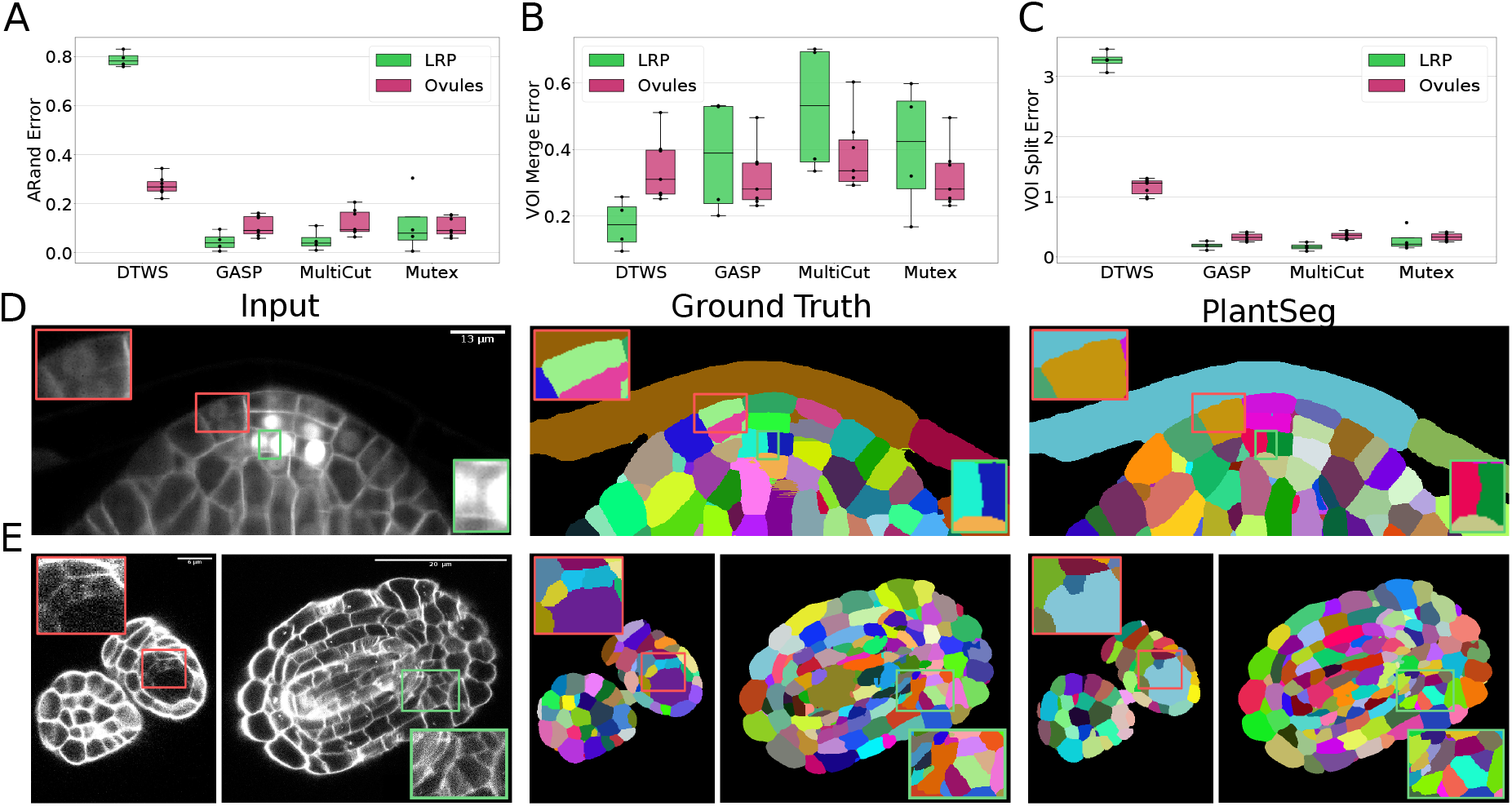
Segmentation using graph partitioning. (A-C) Quantification of segmentation produced by Multicut, GASP, Mutex watershed (Mutex) and DT watershed (DT WS) partitioning strategies. The Adapted Rand error (A) assesses the overall segmentation quality whereas VOI_merge_ (B) and VOI_split_ (C) assess erroneous merge and splitting events. Box plots represent the distribution of values for seven (ovule, magenta) and four (LRP, green) samples. (D, E) Examples of segmentation obtained with PlantSeg on the lateral root (D) and ovule (E) datasets. Green boxes highlight cases where PlantSeg resolves difficult cases whereas red ones highlight errors.

#### 2.1.3 Step 2: segmentation of tissues into cells using graph partitioning

Once the cell boundaries are predicted, segmentation of the cells can be formulated as a generic graph partitioning problem, where the graph is built as the region adjacency graph of the image voxels. However, solving the partitioning problem directly on the voxel-level is computationally very expensive for volumes of biologically relevant size. To make the computation tractable, we first cluster the voxels into so-called supervoxels by running a watershed algorithm on the distance transform of the boundary map, seeded by all local maxima of the distance transform smoothed by a Gaussian blur. The region adjacency graph is then built directly on supervoxels and partitioned into an unknown number of segments to deliver a segmentation. We tested four different partitioning strategies: Multicut [32], hierarchical agglomeration as implemented in GASP average (GASP) [33], Mutex watershed (Mutex) [14] as well as the distance transform (DT) watershed [34] as a baseline since similar methods have been proposed previously [18, 19].

To quantify the accuracy of the four segmentation strategies we use Adapted Rand error (ARand) for the overall segmentation quality and two other metrics based on the variation of information [35], measuring the tendency to over-split (VOI_split_) or over-merge (VOI_merge_), see section 4.4. GASP, Multicut and Mutex watershed consistently produced accurate segmentation on both datasets with low ARand errors and low rates of merge and split errors (Figure 3A-C and Supp. Table 3). As expected DT watershed tends to over-segment with higher split error and resulting higher ARand error. Multicut solves the graph partitioning problem in a globally optimal way and is therefore expected to perform better compared to greedy algorithms such as GASP and Mutex watershed. However, in our case the gain was marginal, probably due to the high quality of the boundary predictions.

The performance of PlantSeg was also assessed qualitatively by expert biologists. The segmentation quality for both datasets is very satisfactory. For example in the lateral root dataset, even in cases where the boundary appeared masked by the high brightness of the nuclear signal, the network correctly detected it and separated the two cells (Figure 3D, green box). On the ovule dataset, the network is able to detect weak boundaries and correctly separate cells in regions where the human expert fails (Figure 3E, green box). The main mode of error identified in the lateral root dataset is due to the ability of the network to remove the nuclear signal which can weaken or remove part of the adjacent boundary signal leading to missing or blurry cell contour. For example, the weak signal of a newly formed cell wall close to two nuclei was not detected by the network and the cells were merged (Figure 3D, red box). For the ovule dataset, in rare cases of very weak boundary signal, failure to correctly separate cells could also be observed (Figure 3E, red box).

Taken together, our data suggest that segmentation using graph partitioning is a good choice for the plant tissue segmentation problem, where boundary discontinuities can sometimes be present.

#### 2.1.4 Performance of PlantSeg on external datasets

To test the generalization capacity of PlantSeg, we assessed its performance on data for which no network training was performed. To this end, we took advantage of the publicly available (https://osf.io/fzr56) Arabidopsis 3D

Digital Tissue Atlas dataset composed of eight confocal stacks of eight different *Arabidopsis thaliana* organs with hand-curated groundtruth [36]. The input images from the digital tissue atlas are similar to the images of the Arabidopsis ovules (confocal stacks of fixed tissue with stained cell contours). We fed the input confocal stacks to PlantSeg and quantitatively and qualitatively assessed the resulting segmentation against the groundtruth. The petal images were not included in our analysis as they are very similar to the leaf and the groundtruth is fragmented, making it difficult to evaluate the results from the pipeline in a reproducible way.

**Table 1:** Quantification of PlantSeg performance on the 3D Digital Tissue Atlas. The Adapted Rand error (ARand) assesses the overall segmentation quality whereas VOI_merge_ and VOI_split_ assess erroneous merge and splitting events.

| Dataset | ARand | VOI<sub>split</sub> | VOI<sub>merge</sub> |
| --- | --- | --- | --- |
| Anther | 0.358 | 0.900 | 0.760 |
| Filament | 0.520 | 1.123 | 1.138 |
| Valve | 0.594 | 0.927 | 1.454 |
| Root | 0.149 | 0.779 | 0.755 |
| Leaf | 0.212 | 0.492 | 0.474 |
| Pedicel | 0.456 | 0.956 | 1.288 |
| Sepal | 0.283 | 0.852 | 1.031 |

Qualitatively, PlantSeg performed well on the 3D Digital Tissue Atlas, giving satisfactory results on all organs, correctly segmenting even the oval non-touching cells of the anther and leaf: a cell shape not present in the training data (Figure 4).

**Figure 4:**
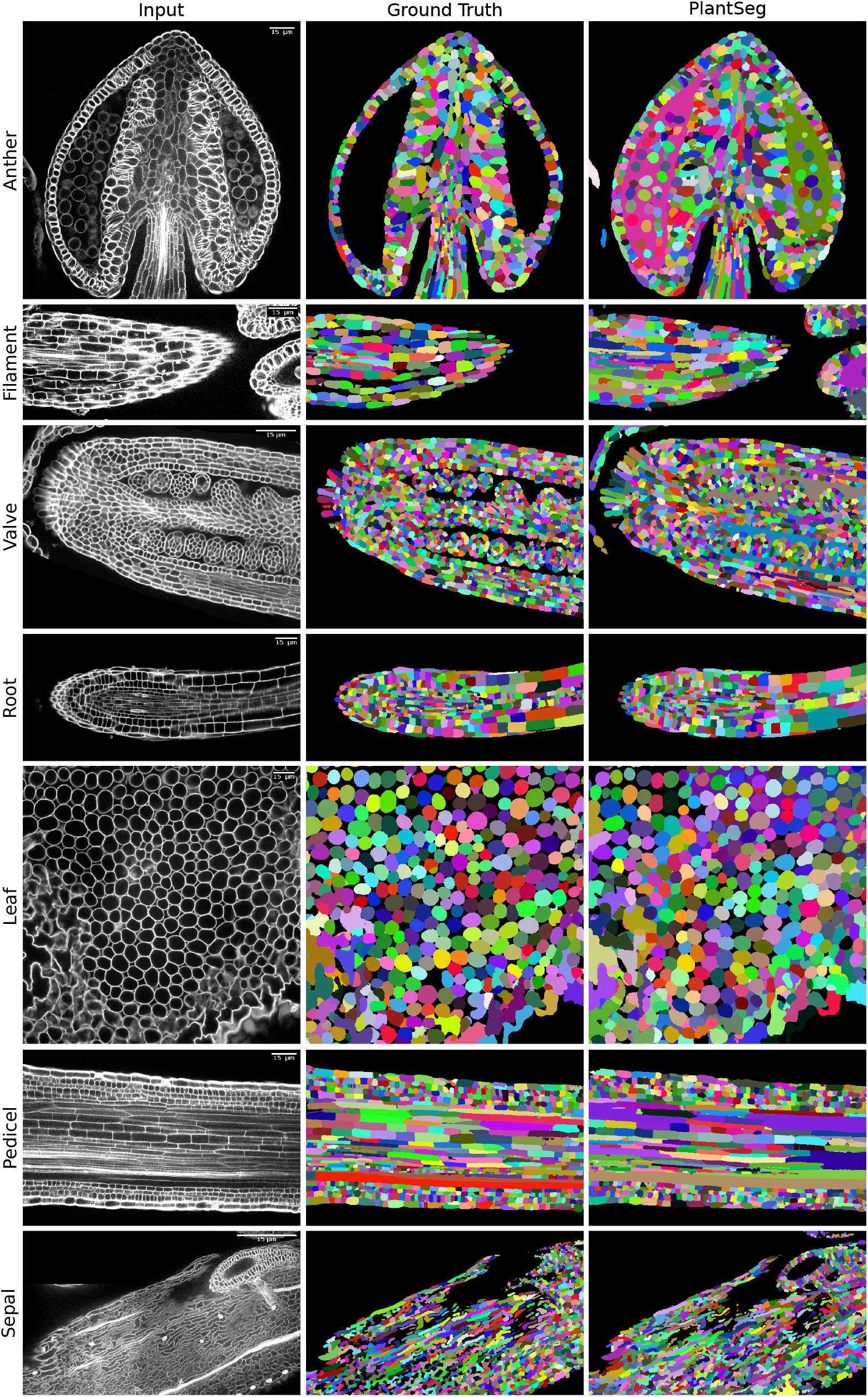
PlantSeg segmentation of different plant organs of the 3D Digital Tissue Atlas dataset, not seen in training. The input image, groundtruth and segmentation results using PlantSeg are presented for each indicated organ.

Quantitatively, the performance of PlantSeg is on par with the scores reported on the LRP and ovules datasets on the anther, leaf, sepal and the root, but lower for the other three tissues (Table 1). However, it should be noted that, the groundtruth included in the dataset was created for analysis of the cellular connectivity network, with portions of the volumes missing or having low quality groundtruth (see e.g filament and sepal in Figure 4). For this reason, the performance of PlantSeg on these datasets may be underestimated.

Altogether, PlantSeg performed well qualitatively and quantitatively on the datasets acquired by a different group, on a different microscope, and at a different resolution than the training data. This demonstrates the generalization capacity of the pre-trained models from the PlantSeg package.

##### Open benchmark performance

For completeness, we have also compared PlantSeg performance with state-of-the-art methods on a non-plant open benchmark consisting of epithelial cells of the *Drosophila* wing disc. Visually, the benchmark images are quite similar to the ovules dataset: membrane staining is used along with a confocal microscope, and the cell shapes are compact and relatively regular. To ensure a fair comparison with the reported benchmark results, we retrain the neural network on the training set provided by the [37], following the same training protocol with a standard 3D U-Net architecture. We found that PlantSeg is very competitive qualitatively and quantitatively (see Supp. Appendix C).

#### 2.1.5 A package for plant tissue segmentation and benchmarking

We release PlantSeg as an open-source software for the 3D segmentation of cells with cell contour staining. PlantSeg allows for data pre-processing, boundary prediction with neural networks and segmentation of the network output with a choice of four partitioning methods: Multicut, GASP, Mutex watershed and DT watershed.

PlantSeg can be executed via a simple graphical user interface (see Figure 5) or via the command line. The critical parameters of the pipeline are specified in a configuration file. Running PlantSeg from the graphical user interface is well suited for processing of single experiments, while the use of the command line utility enables large scale batch processing and remote execution. PlantSeg comes with several networks pre-trained on different voxel size of the Arabidopsis ovule and LRP datasets. Users can select from the available set of pre-trained networks the one with features most similar to their datasets. Alternatively, users can let PlantSeg select the pre-trained network based on the microscope modality (light sheet or confocal) and voxel size. PlantSeg also provides a command-line tool for training the network on new data when none of the pre-trained network is suitable to the user’s needs.

**Figure 5:**
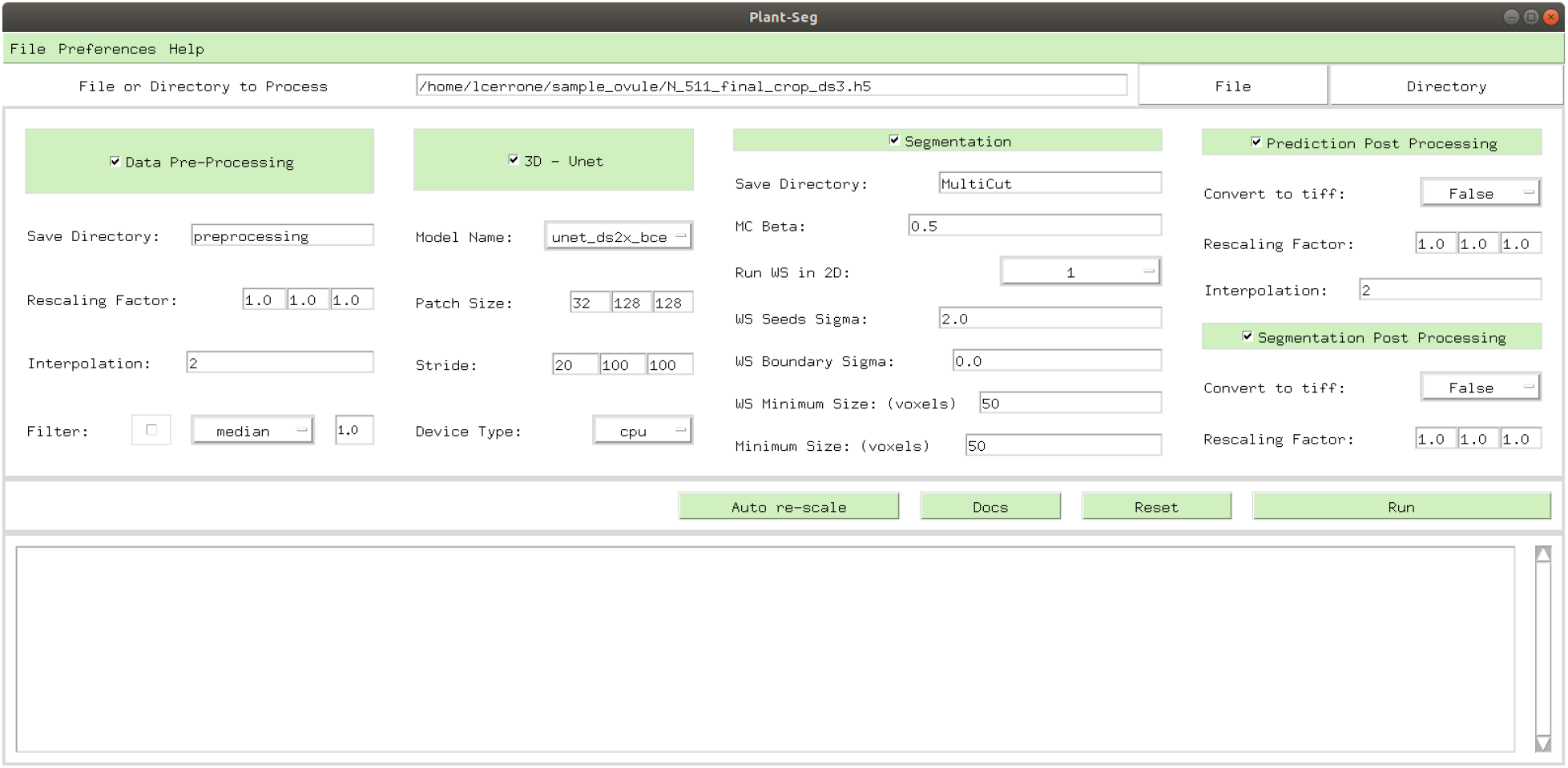
PlantSeg GUI. The interface allows to configure and execute all steps of the segmentation pipeline, such as: selecting the steps to compute, selecting the neural network model to use and specifying hyperparameters of the partitioning algorithm.

PlantSeg is open-source and publicly available https://github.com/hci-unihd/plant-seg. The repository includes a complete user guide, the evaluation scripts used for quantitative analysis, and the employed datasets. Besides the source code, we provide a conda package and a docker image to simplify the installation. The software is written in Python, the neural networks use the PyTorch framework. We also make available the raw microscopic images as well as the groundtruth used for training, validation and testing.

### 2.2 Applications of PlantSeg

#### 2.2.1 Variability in cell number of ovule primordia

Ovule development in *Arabidopsis thaliana* has been described to follow a stereotypical pattern [38, 39]. However, it is unclear if ovules within a pistil develop in a synchronous fashion.

Taking advantage of PlantSeg we undertook an initial assessment of the regularity of primordia formation between ovules developing within a given pistil (Figure 6). We noticed that spacing between primordia is not uniform (Figure 6A). Our results further indicated that six out of the eight analyzed stage 1 primordia (staging according to [39]) showed a comparable number of cells (140.5 10.84, mean SD, ovules 1-5, 7) (Figure 6B). However, two primordia exhibited a smaller number of cells with ovule #6 being composed of 91 and and ovule #8 of 57 cells, respectively. Interestingly, we observed that the cell number of a primordium does not necessarily translate into its respective height or proximal-distal (PD) extension. For example, ovule #2, which is composed of 150 cells and thus of the second largest number of cells of the analyzed specimen, actually represents the second shortest of the eight primordia with a height of 26.5*μm* (Figure 6C). Its comparably large number of cells relates to a wide base of the primordium. Taken together, this analysis indicates that ovule primordium formation within a pistil is relatively uniform, however, some variability can be observed.

**Figure 6:**
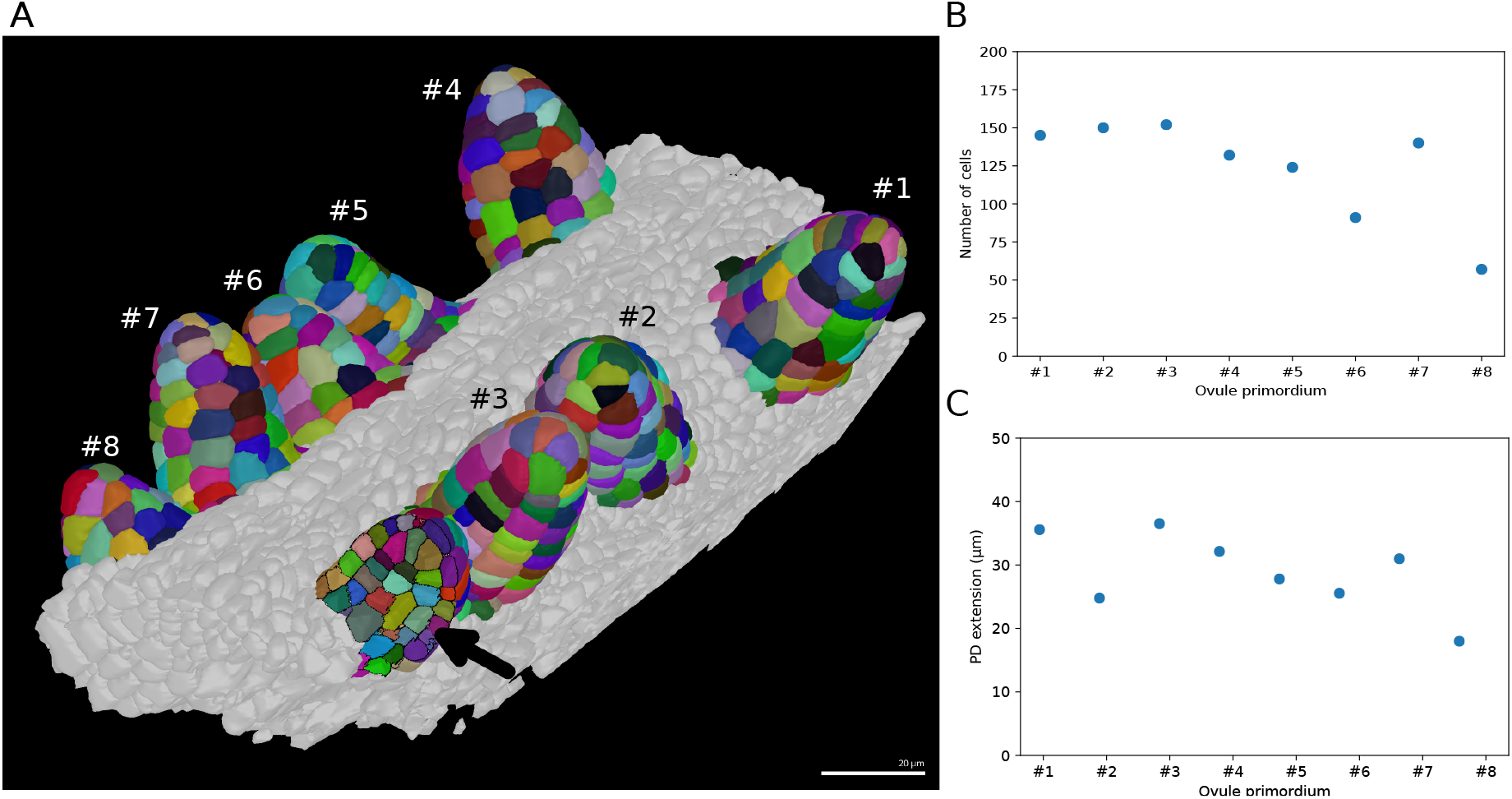
Ovule primordium formation in *Arabidopsis thaliana*. (A) 3D reconstructions of individually labelled stage 1 primordia of the same pistil are shown (stages according to [39]). The arrow indicates an optical mid-section through an unlabeled primordium revealing the internal cellular structure. The raw 3D image data were acquired by confocal laser scanning microscopy according to [26]. Using MorphographX, quantitative analysis was performed on the three-dimensional mesh obtained from the segmented image stack. Cells were manually labelled according to the ovule specimen (from #1 to #8). (B, C) Quantitative analysis of the ovule primordia shown in (A). (B) shows a graph depicting the total number of cells per primordium. (C) shows a graph depicting the proximal-distal (PD) extension of the individual primordia (distance from the base to the tip). Scale bar: 20*μm*.

#### 2.2.2 Asymmetric division of lateral root founder cells

*Arabidopsis thaliana* constantly forms lateral roots (LRs) that branch from the main root. These LRs arise from a handful of founder cells located in the pericycle, a layer of cells located deep within the primary root. Upon lateral root initiation, these founder cells invariably undergo a first asymmetric division giving rise to one small and one large daughter cell. Although the asymmetric nature of this division has long been reported [40, 41] and its importance realised [42], it is still unknown how regular the volume partitioning is between the daughter cells. We used the segmentation of the LR dataset produced by PlantSeg to quantify this ratio. The asymmetric divisions were identified by visual examination during the first 12 hours of the recording and the volumes of the two daughter cells retrieved (Figure 7B). The analysis of cell volume ratios confirmed that the first division of the founder cell is asymmetric with a volume ratio between the two daughter cells of 0.65 *±* 0.22 (mean *±* SD, *n* = 23) (Figure 7C).

**Figure 7:**
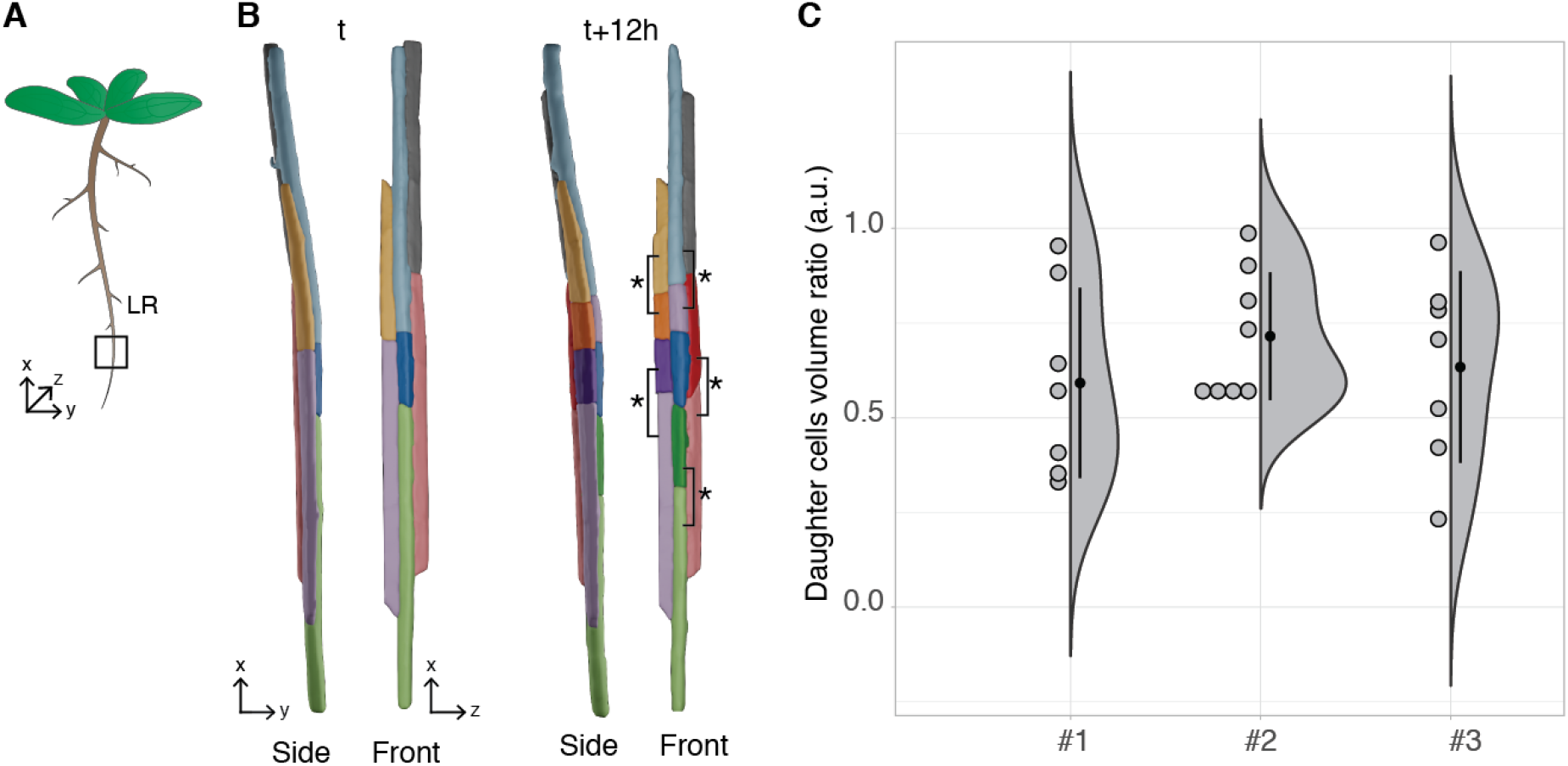
Asymmetric cell division of lateral root founder cells. (A) Schematic representation of *Arabidopsis thaliana* with lateral roots (LR). The box depict the region of the main root that initiates LRs. (B) 3D reconstructions of LR founder cells seen from the side and from the front at the beginning of recording (*t*) and after 12 hours (*t+12*). The star and brackets indicate the two daughter cells resulting from the asymmetric division of a LR founder cell. (C) Half-violin plot of the distribution of the volume ratio between the daughter cells for 3 different movies (#1, #2 and #3). The average ratio of 0.6 indicates that the cells divided asymmetrically.

#### 2.2.3 Epidermal cell volumes in a shoot apical meristem

Epidermal cell morphologies in the shoot apical meristem of *Arabidopsis thaliana* are genetically controlled and even subtle changes can have an impact on organogenesis and pattern formation. To quantify respective cell shapes and volumes in the newly identified big cells in epidermis (*bce*) mutant we used the PlantSeg package to analyze image volumes of six *Arabidopsis thaliana* meristems (three wild type and three *bce* specimens).

Inflorescence meristems of *Arabidopsis thaliana* were imaged using confocal laser scanning microscopy after staining cell walls with DAPI. Image volumes (167 × 167 × 45) *μm* were used to obtain 3D cell segmentations using PlantSeg: in this case a 3D UNet trained on the *Arabidopsis* ovules was used in combination with the Multicut algorithm. This segmentation procedure allows to determine epidermal cell volumes for wild-type (Figure 8A) and the *bce* mutant (Figure 8B). Cells within a radius of 33*μm* around the manually selected center of the meristem (colored cells in Figure 8A and B) were used for the cell volume quantification shown in Figure 8C. The mean volume of epidermal cells in the *bce* mutant is increased by roughly 50 % whereas overall meristem size is only slightly reduced which implicates changes in epidermal cell division in mutant meristems.

**Figure 8:**
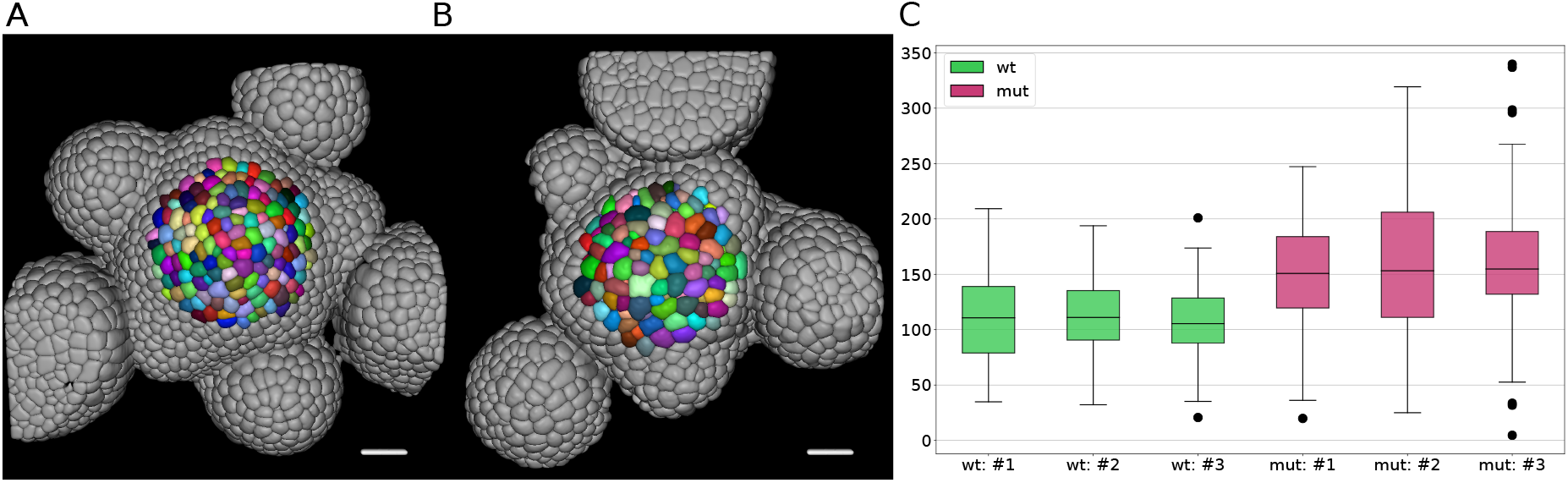
Volume of epidermal cell in the shoot apical meristem of Arabidopsis. Segmentation of epidermal cells in *wildtype* (A) and *bce* mutant (B). Cells located at the center of the meristem are colored. Scale bar: 20*μm*. (C) Quantification of cell volume (*μm*^3^) in 3 different *wildtype* and *bce* mutant specimens.

## 3 Discussion

Taking advantage of the latest developments in machine learning and computer vision we created PlantSeg, a simple, powerful, and versatile tool for plant cell segmentation. Internally, it implements a two-step algorithm: the images are first passed through a state-of-the-art convolutional neural network to detect cell boundaries. In the second step, the detected boundaries are used to over-segment the image using the distance transform watershed and then a region adjacency graph of the image superpixels is constructed. The graph is partitioned to deliver accurate segmentation even for noisy live imaging of dense plant tissue.

PlantSeg was trained on confocal images of *Arabidopsis thaliana* ovules and light sheet images of the lateral root primordia and delivers high-quality segmentation on images from these datasets never seen during training as attested by both qualitative and quantitative benchmarks. We experimented with different U-Net designs and hyperparameters, as well as with different graph partitioning algorithms, to equip PlantSeg with the ones that generalize the best. This is illustrated by the excellent performance of PlantSeg without retraining of the CNN on a variety of plant tissues and organs imaged using confocal microscopy (3D Cell Atlas Dataset). This feature underlines the versatility of our approach for images presenting similar features to the ones trained upon. Of importance, PlantSeg also comes with scripts to train and evaluate the performance of CNN trained de novo on a new set of images. Given the importance of groundtruth for training of CNNs we also provide instructions on how to generate groundtruth (Supp. Appendix A). Besides the plant data, we compared PlantSeg to the state-of-the-art on an open benchmark for the segmentation of epithelial cells in the *Drosophila* wing disc. There, the neural network was re-trained on the benchmark data, but the rest of the pipeline was kept the same as for the plant data. PlantSeg performance was shown to be on par with the benchmark leaders.

We demonstrate the usefulness of PlantSeg on three concrete biological applications that require accurate extraction of cell geometries from complex, densely packed 3D tissues. First, PlantSeg allowed to sample the variability in the development of ovules in a given pistil and reveal that those develop in a relatively synchronous manner. Second, PlantSeg allowed the precise computation of the volumes of the daughter cells resulting from the asymmetric division of the lateral root founder cell. This division results in a large and a small daughter cells with volume ratio of 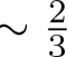 between them. Fninally, segmentation of the epidermal cells in the shoot apical meristem revealed that these cells are enlarged in the *bce* mutant compared to wild type. Accurate and versatile extraction of cell outlines rendered possible by PlantSeg opens the door to rapid and robuts quantitative morphometric analysis of plant cell geometry in complex tissue. This is particularly relevant given the central role plant cell shape plays in the control of cell growth and division [43].

Unlike intensity-based segmentation methods used, for example, to extract DAPI-stained cell nuclei, our approach relies purely on boundary information derived from cell contour detection. While this grants access to the cell morphology and cell-cell interactions, it brings additional challenges to the segmentation problem. Blurry or barely detectable boundaries lead to discontinuities in the membrane structure predicted by the network, which in turn might cause cells to be under-segmented. The segmentation results produced by PlantSeg are not fully perfect and still require proof-reading to reach 100 % accuracy. If nuclei are imaged along with cell contours, nuclear signal can be leveraged for automatic proof-reading as we have explored in [44]. In future work, we envision developing new semi-supervised approaches that would exploit the vast amounts of unlabeled data available in the plant imaging community.

During the development of PlantSeg, we realised that very few benchmark datasets were available to the community for plant cell segmentation tasks, a notable exception being the 3D Tissue Atlas [36]. To address this gap, we publicly release our sets of images and the corresponding hand-curated groundtruth in the hope to catalyse future development and evaluation of cell instance segmentation algorithms. Links to the datasets can be found in the PlantSeg repository at https://github.com/hci-unihd/plant-seg.

## 4 Materials and methods

### 4.1 Biological material and imaging

Imaging of the *Arabidopsis thaliana* ovules was performed as described in [26]. Imaging of the shoot apical meristem was performed as previously described [45, 46] with a confocal laser scanning microscope (Nikon A1, 25× NA=1.1) after staining cell walls with DAPI (0.2mg/ml). For imaging of *Arabidopsis thaliana* lateral root, seedlings of the line sC111 (*UB*10_*pro*_ ∷ *PIP*1,4 − 3 × *GFP/GATA*23_*pro*_ ∷ *H*2*B* : 3 × *mCherry/DR*5*v*2_*pro*_ ∷ 3 × *Y FPnls/RPS*5*A*_*pro*_ ∷ *dtT omato* : *NLS*, described in [27]) were used at 5 day post germination. Sterilized seeds were germinated on top of 4.5 mm long Blaubrand micropipettes (Cat 708744; 100*μ*l) that were immobilised on the bottom of a petri dish and covered with 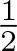 MS-phytagel [47]. Before sowing, the top of the micropipettes is exposed by removing the phytagel with a razor blade and one seed is sowed per micropipette. Plates were stratified for two days and transfered to a growth incubator (23°C, 16h day light). Imaging was performed on a Luxendo MuViSPIM (https://luxendo.eu/products/muvi-spim/) equipped with two 10× NA=0.3 for illumination and 40× NA=0.8 for detection. The following settings were used for imaging: image size 2048 × 2048, exposure time 75ms, channel #1 illumination 488nm 10% power, detection 497-553nm band pass filter, channel #2 illumination 561nm 10 % power, detection 610-628nm band pass filter. Stacks encompassing the whole volume of the root were acquired every 30 minutes. Images from the two cameras were fused using the Luxendo Image processing tool and registered to correct any 3D drift using the BigDataProcessor [48] in Fiji [49].

### 4.2 Neural Network Training and Inference

#### 4.2.1 Training

A 3D U-Net was trained to predict the binary mask of cell boundaries. Groundtruth cell contours where obtained by taking the groundtruth label volume, finding a 2 voxels thick boundaries between labeled regions (*find*_*boundaries*(·) function from the Scikit-image package [50]) and applying a Gaussian blur on the resulting boundary image. Gaussian smoothing reduces the high frequency components in the boundary image, which helps to prevent over-fitting and makes the boundaries thicker, increasing the number of foreground signal during training. Transforming the label image *I_label_*(*x*) into the boundary image *I*_*boundary*_ (*x*) is depicted in Equation 1.

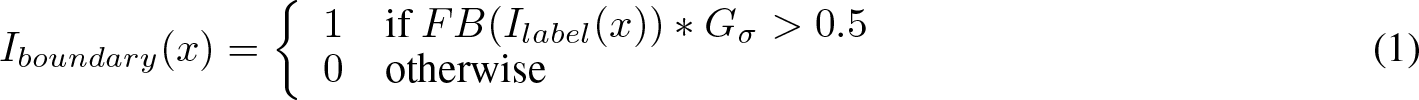

Where *FB*(·) transforms the labeled volume into the boundary image and *G*_*σ*_ is the isotropic Gaussian kernel. We use *σ* = 1.0 in our experiments. The different architectures were trained using Adam optimizer [51] with *β*_1_ = 0.9*, β*_2_ = 0.999, L2 penalty of 0.00001 and initial learning rate *ϵ* = 0.0002. Networks were trained until convergence for 100K iterations, using the PyTorch framework [52] on 8 NVIDIA GeForce RTX 2080 Ti GPUs. For validation during training, we used the adjusted Rand error computed between the groundtruth segmentation and segmentation obtained by thresholding the probability maps predicted by the network and running the connected components algorithm. The learning rate was being reduced by a factor of 2 once the learning stagnated during training, i.e no improvements on the validation set for a given number of iterations. We choose as best network the one with lowest Arand error values. For training with small patch sizes we used 4 patches of shape 100 × 100 × 80 and batch normalization [31] per network iteration. When training with a single large patch (size 170 × 170 × 80), we used group normalization layers [30] instead of batch normalization. This comes from the fact that batch normalization with a single patch per iteration becomes an instance normalization [53] and makes the estimated batch statistics weaker. We also found out that the order of operations at each level of the U-Net impacts the training and the final performance of the network. Normalization layer followed by the 3D convolution and ReLU activation consistently gave better results. We refer to Supp. Table 2 for comparison of different layer ordering. We used random horizontal and vertical flips, random rotations in the XY-plane, elastic deformations [9] and noise augmentations of the input image during training. We found out that noise augmentations in the form of additive Gaussian and additive Poisson noise improve the performance of all tested network variants. The performance of CNNs is sensitive to changes in voxel size and object sizes between training and test images [54]. We thus also trained the networks using version of the dataset down scaled 2× in XY. The 16 networks resulting from the combination of scaling, loss function and architecture can be used by PlantSeg to fit best to the user’s data.

#### 4.2.2 Inference

During inference we used mirror padding on the input image to improve the prediction at the boundaries of the volume. We kept the same patch sizes as during training since increasing it during inference might lead to decreased quality of the predictions, especially on the borders of the patch. We also parse the volume patch-by-patch with a 50 % overlap between consecutive tiles and average the probability maps. This strategy prevents checkerboard artifacts and reduces noise in the final prediction.

The code used for training and inference can be found at https://github.com/hci-unihd/pytorch-3dunet.

### 4.3 Segmentation using graph partitioning

The boundary predictions produced the CNN are treated as a graph *G*(*V, E*), where nodes *V* are represented by the image voxels, and the edges *E* connect adjacent voxel. The weight *w* ∈ *R*^+^ of each edge, is derived from the probability maps. On this graph we first performed an over-segmentation by running the DT watershed [34]. For this, we threshold the boundary probability maps at a given value (0.4 in our case). Then we compute the distance transform from the binary boundary image, apply a Gaussian smoothing (*sigma* = 2.0) and assign a seed to every local minimum in the resulting distance transform map. Finally we removed regions small regions (< 50 voxels). Standalone DT watershed already delivers accurate segmentation and can be used as is in simple cases when for example noise is low and/or boundaries are sharp.

For Multicut [32], GASP [33], and Mutex watershed [14] algorithms, we used the DT watershed as an input. Although all three algorithms could be run directly on the boundary predictions produced the CNN (voxel level), we choose to run them on a region adjacency graph (RAG) derived from the DT watershed to reduce the computation time. In the region adjacency graph each node represents a region and edges connect adjacent regions. We compute edge weights by using the mean value of the probabilities maps along the boundary. We then run Multicut, GASP or Mutex watershed with a hyperparameter *beta* = 0.6 that balances over-and under-segmentation (with higher *β* tending to over-segment).

### 4.4 Metrics used for evaluation

For the boundary predictions we used precision (number of true positive divided by the number of positive observation), recall (number of true positive divided by the sum of true positive and false positive) and F1 score

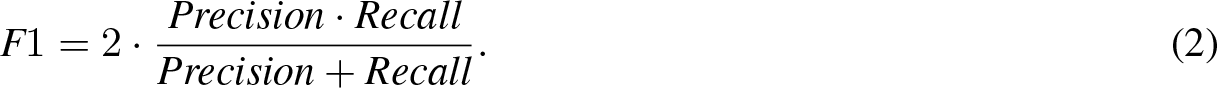

For the final segmentation, we used the inverse of the Adjusted Rand Index (AdjRand) [55] defined as ARand error = 1 − AdjRand [56] which measures the distance between two clustering as global measure of accuracy between PlantSeg prediction and groundtruth. A ARand error of 0 means that the PlantSeg results are identical to the groundtruth, whereas 1 shows no correlation. To quantify the rate of merge and split errors, we used the Variation of Information (VOI) which is an entropy based measure of clustering quality [57]. It is defined as:

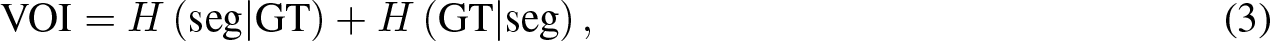

where *H* is the conditional entropy function and the Seg and GT the predicted segmentation and groundtruth segmentation respectively. *H* (seg|GT) defines the split mistakes (VOI_split_) whereas *H* (GT|Seg) is the merge mistake (VOI_merge_).

## 5 Authors contribution

AW, LC, CP, AB, FAH, AM and AK designed and implemented PlantSeg. AV, RT, ML, AVB, SS, SD-N, GWB, KS and AM contributed the datasets, groundtruth and analysis of the ovule and lateral root datasets. CW and JL provided analysis of the shoot meristem. AW, LC, KS, FAH, AM and AK wrote the manuscript.

## 6 Declaration of interests

The authors declare no competing interests.

## 7 Acknowledgments

We thank Kerem Celikay, Melanie Holzapfel and Boyko Vodenicharski for their help in the early steps of the project. We further acknowledge the support of the Center for Advanced Light Microscopy (CALM) at the TUM School of Life Sciences. GWB and SD-N were funded by Leverhulme Grant RPG-2016-049. CP and AK were funded by Baden-Wuerttemberg Stiftung. This work was supported by the DFG FOR2581 to the Hamprecht (P3), Kreshuk (P3), Lohmann (P5), Maizel (P6) and Schneitz (P7) labs.

## SUPPLEMENTARY MATERIAL

### A Groundtruth Creation

Training state-of-the-art deep neural network for semantic segmentation requires large amount of densely annotated samples. Groundtruth creation for the ovule dataset has been described previously [1], we briefly describe here how the dense groundtruth labeling of cells for the lateral root was generated.

We bootstrapped the process using the Autocontext Workflow [2] of the open-source ilastik software [3] which is used to segment cell boundaries from sparse user input (scribbles). It is followed by the ilastik’s multicut workflow [4] which takes the boundary segmentation image and produces the cell instance segmentation. These initial segmentation results were iteratively refined (see Figure 1). First, the segmentation is manually proofread in a few selected regions of interest using the open-source Paintera software [5]. Second, a state-of-the-art neural network is trained for boundary detection on the manually corrected regions. Third, PlantSeg framework consisting of neural network prediction and image partitioning algorithm is applied to the entire dataset resulting in a more refined segmentation. The 3-step iterative process was repeated until an instance segmentation of satisfactory quality was reached. A final round of manual proofreading with Paintera is performed to finalize the groundtruth creation process.

### B Exploiting nuclei staining to improve the lateral root cells segmentation

The lateral root dataset contains a nuclei marker in a separate channel. In such cases we can take advantage of the fact that a cell contains only one nuclei and use this information as an additional clue during segmentation.

For this, we first segmented the nuclei using a simple and accurate segmentation pipeline that consists of a 3D U-Net trained to predict the binary nuclei mask followed by thresholding of the probability maps and connected components. We then incorporated this additional information into the multicut formulation, called lifted multicut [6], where additional repulsive edges are introduced between the nodes in the graph corresponding to the different nuclei segmented from the second channel.

We compared the scores of this lifted multicut algorithm to the scores for GASP, multicut and mutext watershed (see Table 3). We see that lifted multicut outperforms not only the standard multicut, but also all the other algorithms. This is because lifted multicut is able to separate two cells incorrectly merged into one region by the segmentation algorithm, as long as the region contains the two different nuclei instances corresponding to the merged cells.

**Supp. Figure 1:**
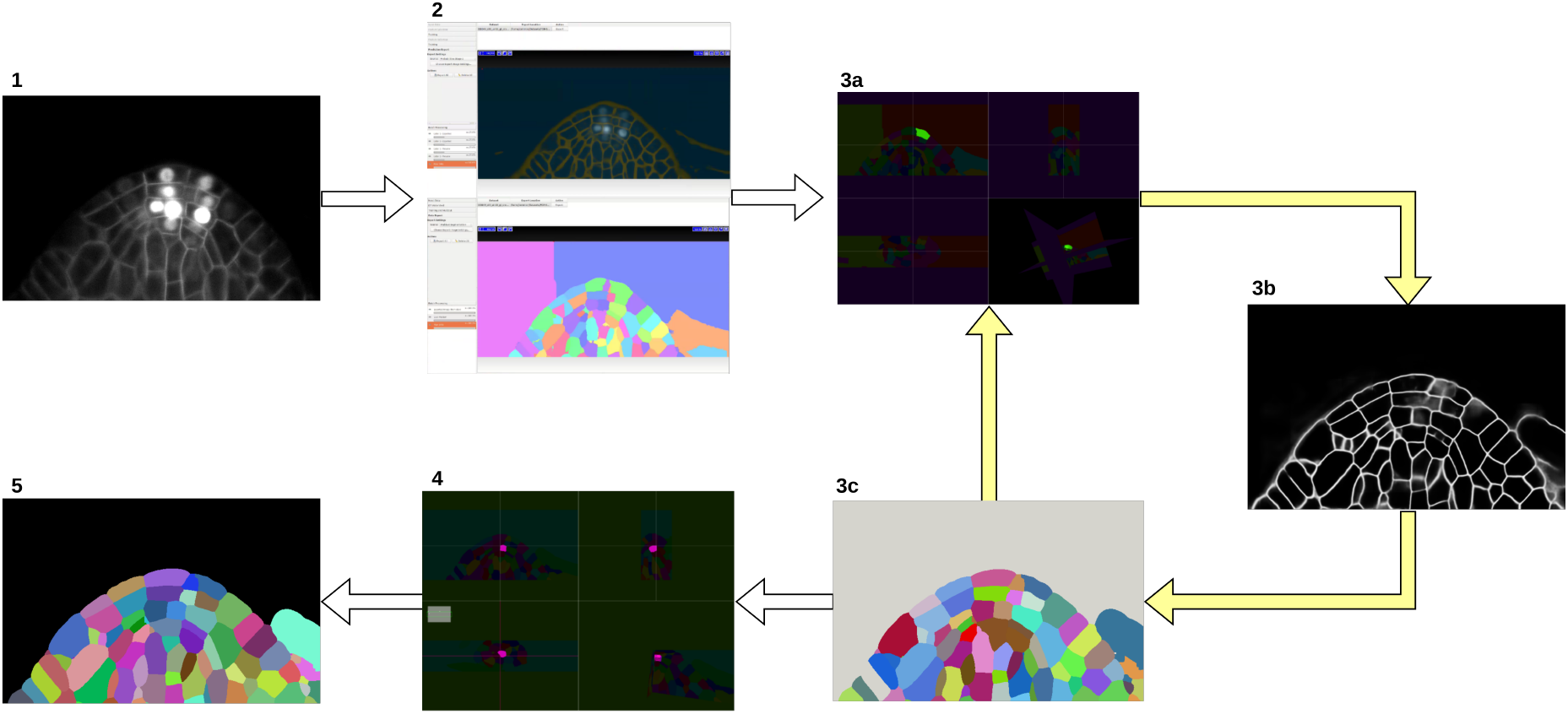
Groundtruth creation process. Starting from the input image (1), an initial segmentation is obtained using ilastik Autocontext followed by the ilastik multicut workflow (2). Paintera is used to proofread the segmentation (3a) which is used for training a 3D UNet for boundary detection (3b). A graph partitioning algorithm is used to segment the volume (3c). Steps 3a, 3b and 3c are iterated until a final round of proofreading with Paintera (4) and the generation of satisfactory final groundtruth labels (5) is obtained.

**Supp. Table 1:** Epithelial Cell Benchmark results. We compare PlantSeg to four other methods using the standard SEG metric [14] (higher is better). Mean and standard deviation of the SEG score are reported for peripodial (3 movies) and proper disc (5 movies) cells.

| Method | PERIPODIAL | PROPER DISC |
| --- | --- | --- |
| MALA | $0.907 \pm 0.029$ | $0.817 \pm 0.009$ |
| FFN | $0.879 \pm 0.035$ | $0.796 \pm 0.013$ |
| MLT-GLA | $0.904 \pm 0.026$ | $0.818 \pm 0.010$ |
| TA | - | $0.758 \pm 0.009$ |
| PlantSeg (multicut) | $0.885 \pm 0.056$ | $0.800 \pm 0.015$ |

### C Performance of PlantSeg on an independent reference benchmark

To test how versatile our approach is, we assessed PlantSeg performance on a non-plant dataset consisting of 2D+t videos of membrane-stained developing *Drosophila* epithelial cells [7]. The benchmark dataset and the results of four state-of-the-art segmentation pipelines are reported in [8]. Treating the movie sequence as 3D volumetric images not only resembles the plant cell images shown in our study, but also allows to pose the 2D+t segmentation as a standard 3D segmentation problem. For a fair comparison of our pipeline with the methods described in [8] we trained a standard 3D U-Net on the epithelial cell dataset. We used the same patch sizes, number of iterations and optimizer hyperparameters as described for the plant samples. For segmentation, we only used the multicut pipeline.

We compared the performance of PlantSeg on the 8 movies of this dataset to the four reported pipelines: MALA [9], Flood Filling Networks (FFN) [10], Moral Lineage Tracing (MLT) [11, 12] and Tissue Analyzer (TA) [8].

Qualitative results of our pipeline are shown in Figure 2: PlantSeg gives close to optimal segmentation results on both the peripodial and proper imaginal disc cells. This impression is confirmed by quantitative benchmark results (see Table 1). Looking at the average run-times of the methods reported in the benchmark shows that the efficient implementation of the multicut solver [13, 4] used in our pipeline is clearly the fastest approach with the average run-time of 1.5 min when run on a modern laptop versus 35 min (MALA), 42 min (MLT) and 90 min (FFN).

Thus, PlantSeg achieves competitive results on-par with other proven methods in terms of accuracy, ranking 3rd behind the MALA and MLT-GLA and outperforming all methods in term of computing time.

**Supp. Figure 2:**
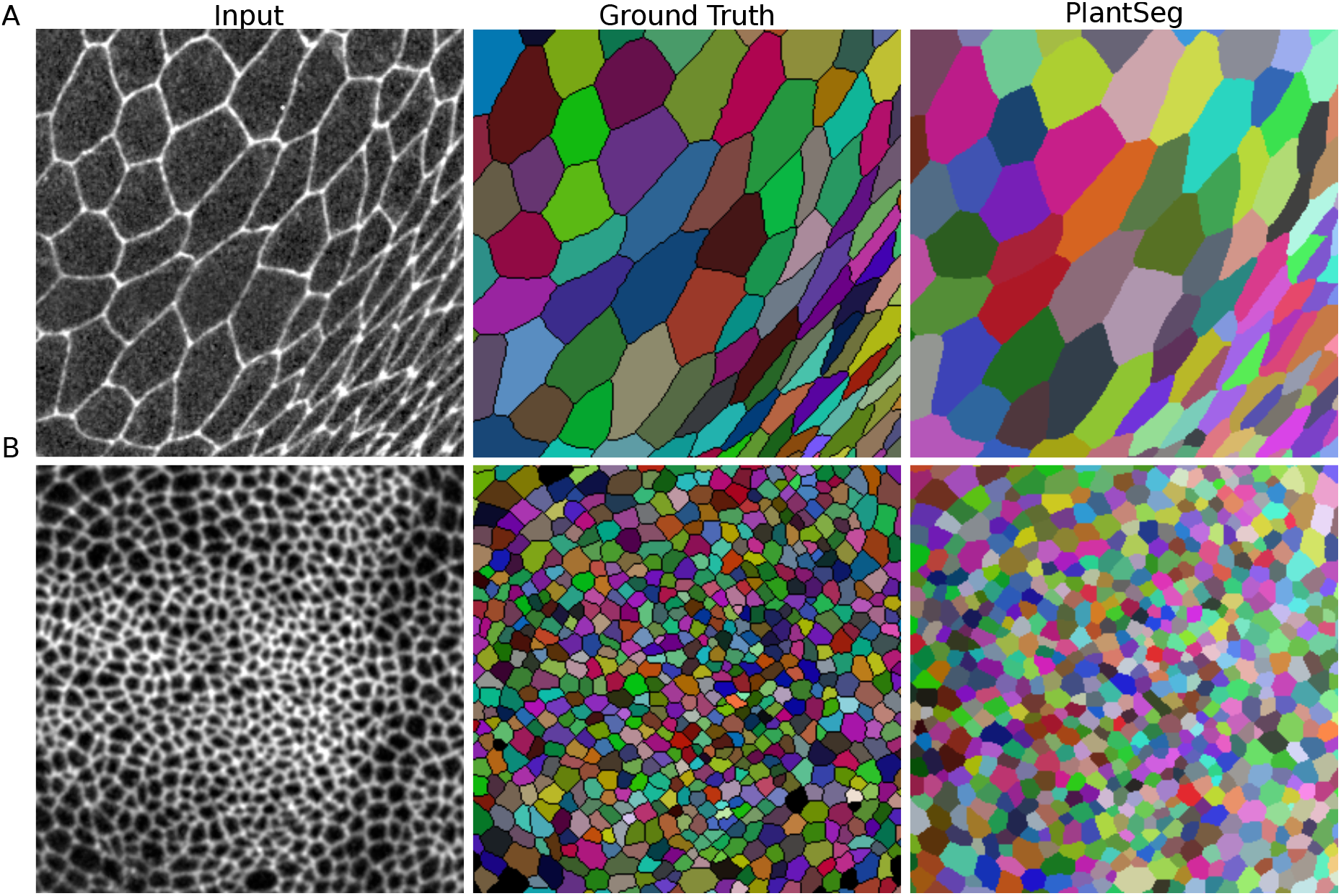
Qualitative results on the Epithelial Cell Benchmark. From top to bottom: Peripodial cells (A), Proper disc cells (B). From left to right: raw data, groundtruth segmentation, multicut segmentation.

### D Supplemental Figures

**Supp. Figure 3:**
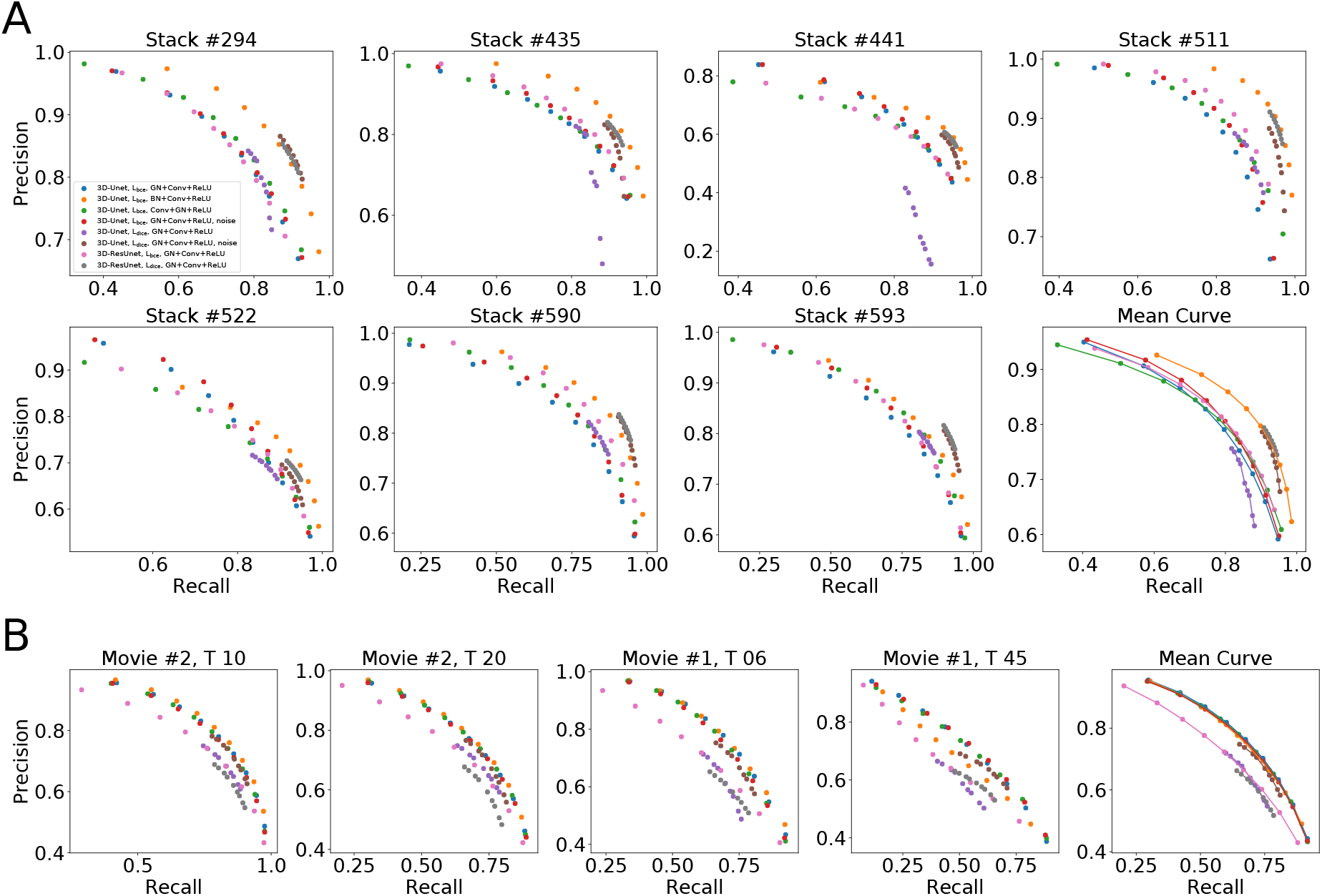
Precision-recall curves on individual stacks for different CNN variants on the ovule (A) and lateral root primordia (B) datasets. Efficiency of boundary prediction was assessed for seven training procedures that sample different type of architecture (3D U-Net *vs.* 3D Residual U-Net), loss function (binary cross-entropy (bce) *vs.* Dice), normalization (Group-Norm (GN) *vs.* Batch-Norm (BN)) and noise augmentation. The larger the area under the curve, the better the precision.

### E Supplemental Tables

**Supp. Table 2:**
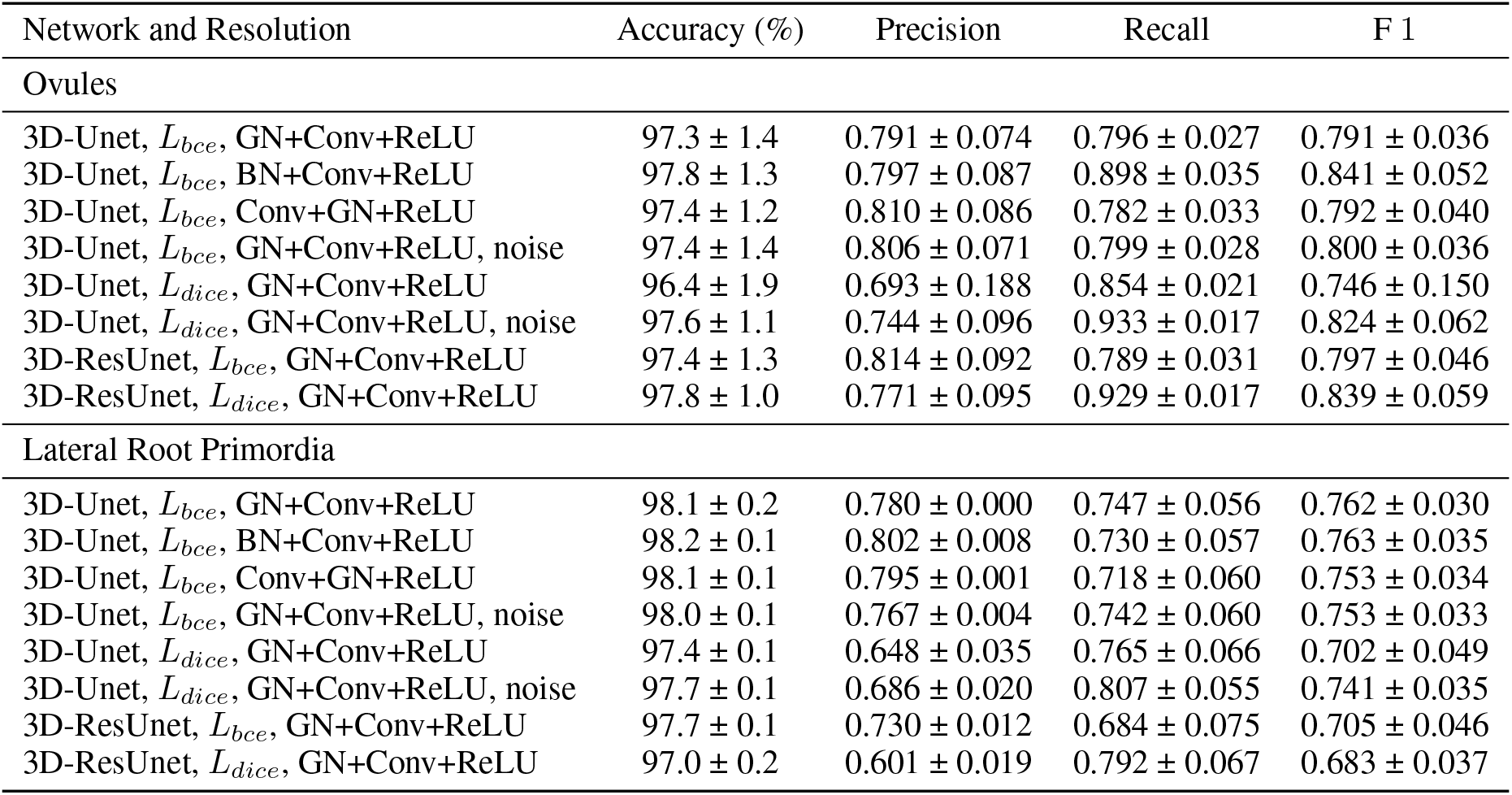
Ablation study of boundary detection accuracy. Efficiency of boundary prediction was assessed for eight training procedures that sample different type of architecture (3D U-Net vs. 3D Residual U-Net), loss function (binary cross-entropy vs. Dice), normalization (Group-Norm vs. Batch-Norm), layer ordering and noise augmentation. All entries are evaluated at a fix threshold of 0.5. Values are obtained from a set of seven specimen for the ovules and four for the lateral root primordia, while the error is measured by standard deviation.

**Supp. Table 3:**
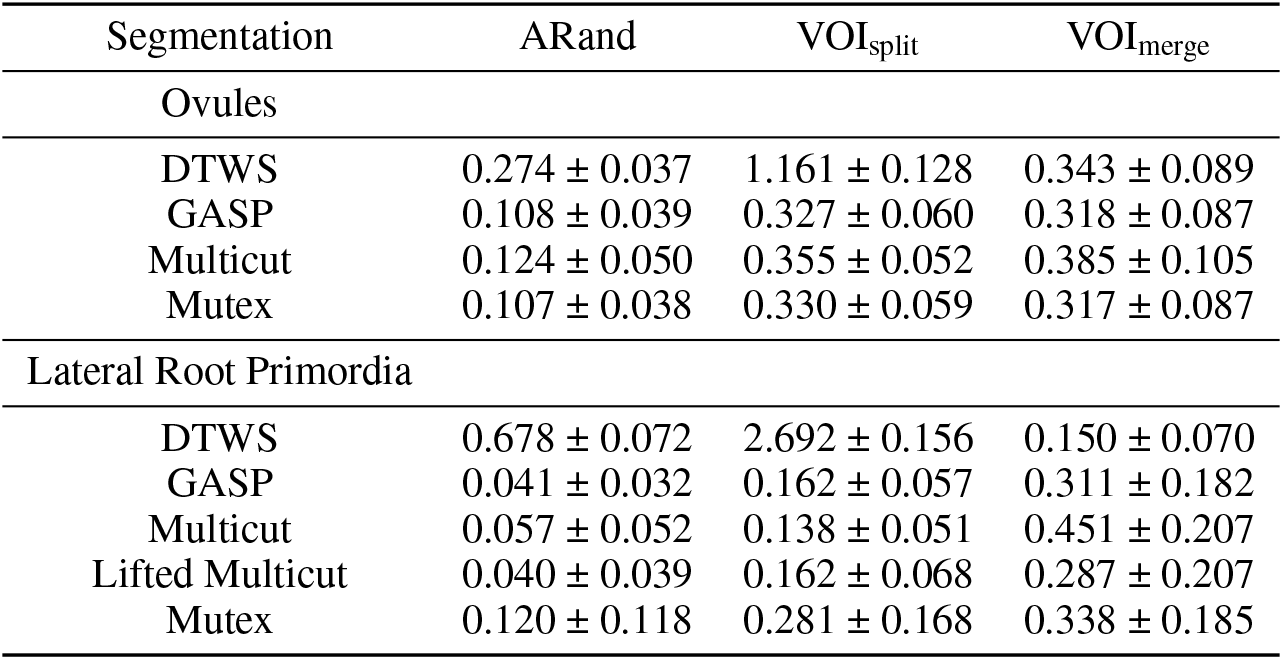
Average segmentation accuracy for different segmentation algorithms. The average is computed from a set of seven specimen for the ovules and four for the lateral root primordia (LRP), while the error is measured by standard deviation. The segmentation is produced by multicut, GASP, mutex watershed (Mutex) and DT watershed (DTWS) clustering strategies. We additionally report the scores given by the lifted multicut on the LRP dataset. The Metrics used are the Adapted Rand error to asses the overall segmentation quality, the VOI_merge_ and VOI_split_ respectively assessing erroneous merge and splitting events (lower is better for all metrics).

| Segmentation | ARand | VOI<sub>split</sub> | VOI<sub>merge</sub> |
| --- | --- | --- | --- |
| Ovules |  |  |  |
| DTWS | 0.274 ± 0.037 | 1.161 ± 0.128 | 0.343 ± 0.089 |
| GASP | 0.108 ± 0.039 | 0.327 ± 0.060 | 0.318 ± 0.087 |
| Multicut | 0.124 ± 0.050 | 0.355 ± 0.052 | 0.385 ± 0.105 |
| Mutex | 0.107 ± 0.038 | 0.330 ± 0.059 | 0.317 ± 0.087 |
| Lateral Root Primordia |  |  |  |
| DTWS | 0.678 ± 0.072 | 2.692 ± 0.156 | 0.150 ± 0.070 |
| GASP | 0.041 ± 0.032 | 0.162 ± 0.057 | 0.311 ± 0.182 |
| Multicut | 0.057 ± 0.052 | 0.138 ± 0.051 | 0.451 ± 0.207 |
| Lifted Multicut | 0.040 ± 0.039 | 0.162 ± 0.068 | 0.287 ± 0.207 |
| Mutex | 0.120 ± 0.118 | 0.281 ± 0.168 | 0.338 ± 0.185 |

